# IMPDH Cytoophidia in the Pancreas: Morphological Dynamics and Metabolic Implications

**DOI:** 10.1101/2025.02.20.639403

**Authors:** Yujie Huang, Ji-Long Liu, Wenchuan Zhang

## Abstract

Cytoophidia are filamentous structures formed by metabolic enzymes, playing critical roles in cellular metabolism and regulation. Inosine monophosphate dehydrogenase (IMPDH), the rate-limiting enzyme in de novo nucleotide synthesis, is known to form cytoophidia and is implicated in cell proliferation, nutrient sensing, and metabolic stress responses. Despite their significance, the behavior of IMPDH cytoophidia under different metabolic conditions, particularly in diabetes, remains poorly understood. This study investigates the distribution, morphology, and assembly patterns of IMPDH cytoophidia in the pancreas of normal and diabetic models. Using immunofluorescence staining, pharmacological interventions (e.g., mycophenolic acid), and glucose tolerance tests, we demonstrate that IMPDH cytoophidia exhibit distinct morphological changes in pancreatic islets under diabetic conditions. Notably, cytoophidia elongation is associated with impaired glucose tolerance, suggesting a potential link between cytoophidia dynamics and metabolic dysfunction. Our findings provide novel insights into the role of IMPDH cytoophidia in metabolic regulation and highlight their potential as therapeutic targets for diabetes and related metabolic disorders.

## INTRODUCTION

Cytoophidia are filamentous structures formed by metabolic enzymes(1), including inosine monophosphate dehydrogenase (IMPDH) (2), the rate-limiting enzyme in the de novo nucleotide synthesis pathway. IMPDH catalyzes the conversion of inosine monophosphate (IMP) to xanthosine monophosphate (XMP) (3) and has been observed to form cytoophidia in various tissues(4; 5), including the mouse pancreas(6) and human cells(7). However, the presence and role of IMPDH cytoophidia in the rat pancreas remain unexplored, representing a critical gap in our understanding of these structures.

The assembly of IMPDH cytoophidia is regulated by multiple factors, including CTP pool size(8), macromolecular crowding(9), and interactions with GTP/GDP(10–12). The inhibitor of IMPDH, such as Mycophenolic acid (MPA) and Ribavirin(13; 14), has been shown to induce cytoophidia formation both in vitro and in vivo(15; 16). These structures are implicated in immune cell proliferation(17–19), cellular nutrient sensing(20), and metabolic regulation(21; 22), highlighting their broad functional significance.

The pancreas, a central metabolic organ, comprises exocrine and endocrine components(23). The endocrine pancreas, particularly the islets of Langerhans, plays a critical role in glucose homeostasis through the secretion of insulin and glucagon. Dysregulation of pancreatic function is a hallmark of metabolic disorders such as diabetes(24), which is characterized by elevated free fatty acids (FFAs) (25), including oleic acid(26). Notably, oleic acid has been shown to promote IMPDH translocation to lipid bodies(27), suggesting a potential link between lipid metabolism and cytoophidia dynamics. Previous studies have also demonstrated that IMPDH cytoophidia colocalize with insulin granules in mouse pancreas(28), and their abundance decreases during fasting(20), further implicating these structures in metabolic regulation.

Despite these advances, the specific functions of IMPDH cytoophidia in pancreatic cells, particularly under diabetic conditions, remain poorly understood. Moreover, research has largely been confined to mouse models, limiting the generalizability of findings. To address these gaps, we investigated the distribution, morphology, and assembly of IMPDH cytoophidia in the pancreas of both normal and diabetic models. Using immunofluorescence staining, pharmacological interventions (e.g., MPA), and metabolic assays, we explored the relationship between cytoophidia dynamics and glucose tolerance. Our findings provide new insights into the role of IMPDH cytoophidia in metabolic regulation and offer potential avenues for therapeutic interventions in metabolic diseases such as diabetes.

## MATERIALS AND METHODS

### Animals

All animal experiments were approved by the Shanghai Jiao Tong University Animal Care and Use Committee and the ShanghaiTech University Experimental Animal Care Committee, and conducted in accordance with the guidelines established by the National Health and Family Planning Commission of China. Animals were housed under specific pathogen-free conditions with free access to water and standard chow.

- Eight-week-old male C57BL/6JGpt mice (Strain NO. N000013) and BKS-db mice (Strain NO. T002407) were purchased from GemPharmatech (Nanjing, China).

- Male Sprague-Dawley rats (200–240 g) were obtained from the Shanghai Lab Animal Research Center.

### Diabetic Models

Streptozotocin (STZ)-induced diabetic models were established in male Sprague-Dawley rats (200–220 g) (29). The experimental group received an intraperitoneal injection of 60 mg/kg STZ (S8050, Solarbio) dissolved in 1% citrate buffer (pH 4.5), while the control group received citrate buffer alone. Three days post-injection, fasting blood glucose levels were measured, and a glucose concentration exceeding 11.1 mmol/L was used as the criterion for successful diabetes induction.

### Frozen Sections

Pancreatic tissues were fixed in 4% paraformaldehyde overnight at 4 °C, followed by stepwise incubation in 15% to 30% sucrose (S1888, Sigma) until the tissues sank. Tissues were embedded in OCT mounting medium (4583, Tissue-Tek) and frozen at −80 °C. Sections (10 μm thick) were cut using a ThermoFisher NX50 cryostat.

### Immunofluorescence

Frozen pancreatic sections were washed with PBS to remove OCT compound, blocked for 2 hours in blocking buffer (2.5% bovine serum albumin, 0.2% Triton X-100, and 1× PBS), and incubated with primary antibodies overnight at room temperature. After washing, sections were incubated with secondary antibodies and Hoechst 33342 (1:10,000, 62249, Thermo) overnight, washed again, and mounted in 50% glycerol.

Primary antibodies:

- Mouse anti-insulin (1:500, 66198-1-IG, Proteintech)

- Rabbit anti-IMPDH2 (1:500, 12948-1-AP, Proteintech) Secondary antibodies:

- Alexa-Fluor-488-conjugated goat anti-mouse IgG (1:500, A11029, Invitrogen)

- Alexa-Fluor-546-conjugated goat anti-rabbit IgG (1:500, A11035, Invitrogen)

- Alexa-Fluor-594-conjugated goat anti-mouse IgG (1:500, A11032, Invitrogen)

### Histology Staining

Pancreatic tissues were fixed in 4% paraformaldehyde, dehydrated, embedded in paraffin, and sectioned. Sections were dewaxed in xylene, rehydrated in gradient ethanol, and stained with hematoxylin and eosin (H&E).

### Pancreatic Islet Isolation and Culture

Islets were isolated from 8-week-old C57BL/6J mice(30). The pancreas was inflated via the bile duct with 0.15 mg/mL collagenase P in Hank’s Balanced Salt Solution, excised, and digested at 37 °C for 15 minutes. Digested tissue was filtered through a 400-mesh sieve and a 70 μm filter, and intact islets were handpicked under a microscope. Islets were cultured in RPMI-1640 medium supplemented with 10% FBS, 7 mM glucose, 100 U/mL penicillin, and 100 μg/mL streptomycin at 37 °C in a humidified atmosphere of 95% O2 and 5% CO2.

### Intraperitoneal Glucose Tolerance Test (IPGTT)

Mice were fasted for 16 hours and injected intraperitoneally with 2 g/kg glucose(31). Blood glucose levels were measured at indicated time points, and the area under the curve (AUC) was calculated after subtracting baseline values.

### Microscopy

Images were acquired using a 63× objective on laser scanning confocal microscopes (Zeiss 880 and Leica SP8). Z-stack images were captured at optimal intervals for quantitative analysis.

### Data Analysis

Confocal images were processed using ZEN 3.8 and ImageJ. Quantitative data were analyzed using Excel and GraphPad Prism 8. Statistical significance was determined using Student’s t-test.

Data were obtained from at least three independent experiments, with more than 50 islets quantified per repeat.

## RESULTS

### IMPDH Forms Cytoophidia in Mouse and Rat Pancreas

The pancreas comprises endocrine and exocrine regions, with the endocrine portion represented by the islets of Langerhans. These islets contain hormone-secreting cells, including α-cells (glucagon) and β-cells (insulin). To investigate the presence of IMPDH- associated cytoophidia in pancreatic tissues, we collected pancreatic samples from wild-type C57BL/6J mice and Sprague-Dawley (SD) rats and prepared them as frozen sections. Immunostaining was performed using antibodies against IMPDH (to label cytoophidia), insulin (to label islets), and Hoechst 33342 (to label DNA). Our results confirmed the presence of IMPDH cytoophidia in both mouse and rat pancreatic tissues (Fig. 1A-C).

**Figure 1.**
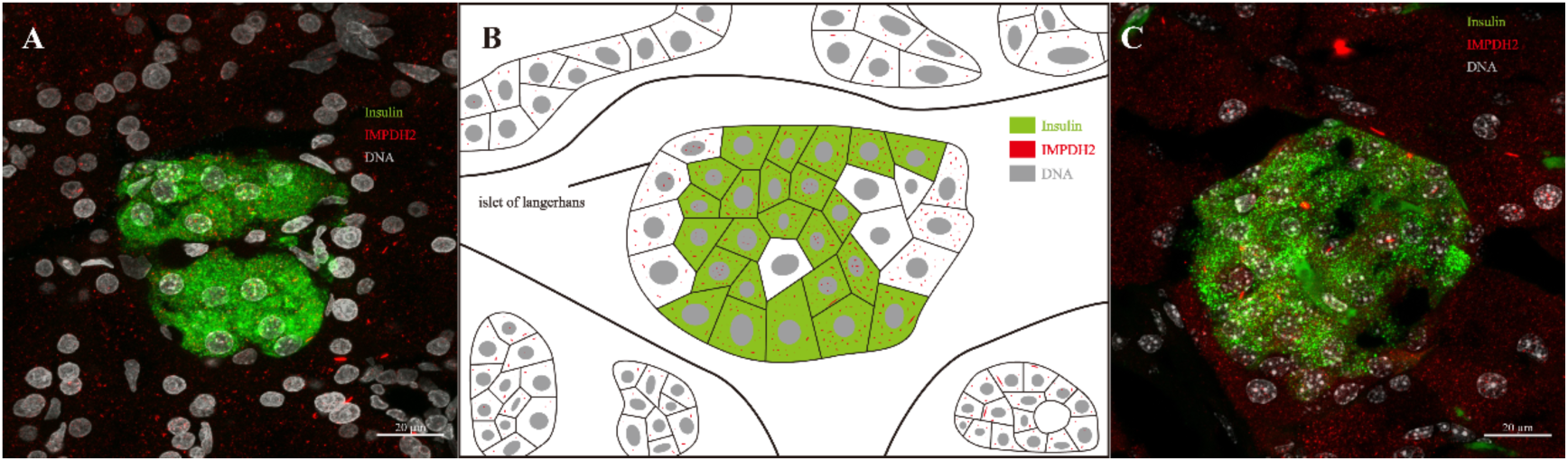
IMPDH forms cytoophidia in the pancreas of mice and rats. **(A)** Immunofluorescence staining of a normal SD rat pancreas section showing IMPDH cytoophidia. Scale bar: 20 μm. **(B)** Schematic illustration of IMPDH-associated cytoophidia in the pancreas, based on panels A and C. **(C)** Immunofluorescence staining of a normal C57BL/6J mouse pancreas section showing IMPDH cytoophidia. Scale bar: 20 μm. In (A-C), IMPDH is shown in red, insulin in green, and Hoechst 33342 (DNA, nuclei) in light gray.

### Distribution of IMPDH Cytoophidia in Endocrine versus Exocrine Pancreas

To further characterize the distribution of IMPDH cytoophidia within the pancreas, we divided the tissue into islet (endocrine) and non-islet (exocrine) regions for quantitative analysis (Fig. 2A-D). Statistical analysis revealed a significantly higher abundance of IMPDH cytoophidia within the islets compared to the surrounding exocrine tissue (Fig. 2F, H). Additionally, we identified two distinct morphological types of cytoophidia: shorter, thinner structures and longer, thicker structures (indicated by yellow arrows in Fig. 2A-D). Notably, shorter cytoophidia were predominantly localized within the islets, whereas longer cytoophidia were more frequently observed in the exocrine regions. To quantify these differences, we measured the area of individual cytoophidia, ranked them by size, and analyzed the top three in each image (Fig. 2E, F). This analysis highlighted significant regional variations in cytoophidium morphology and distribution.

**Figure 2.**
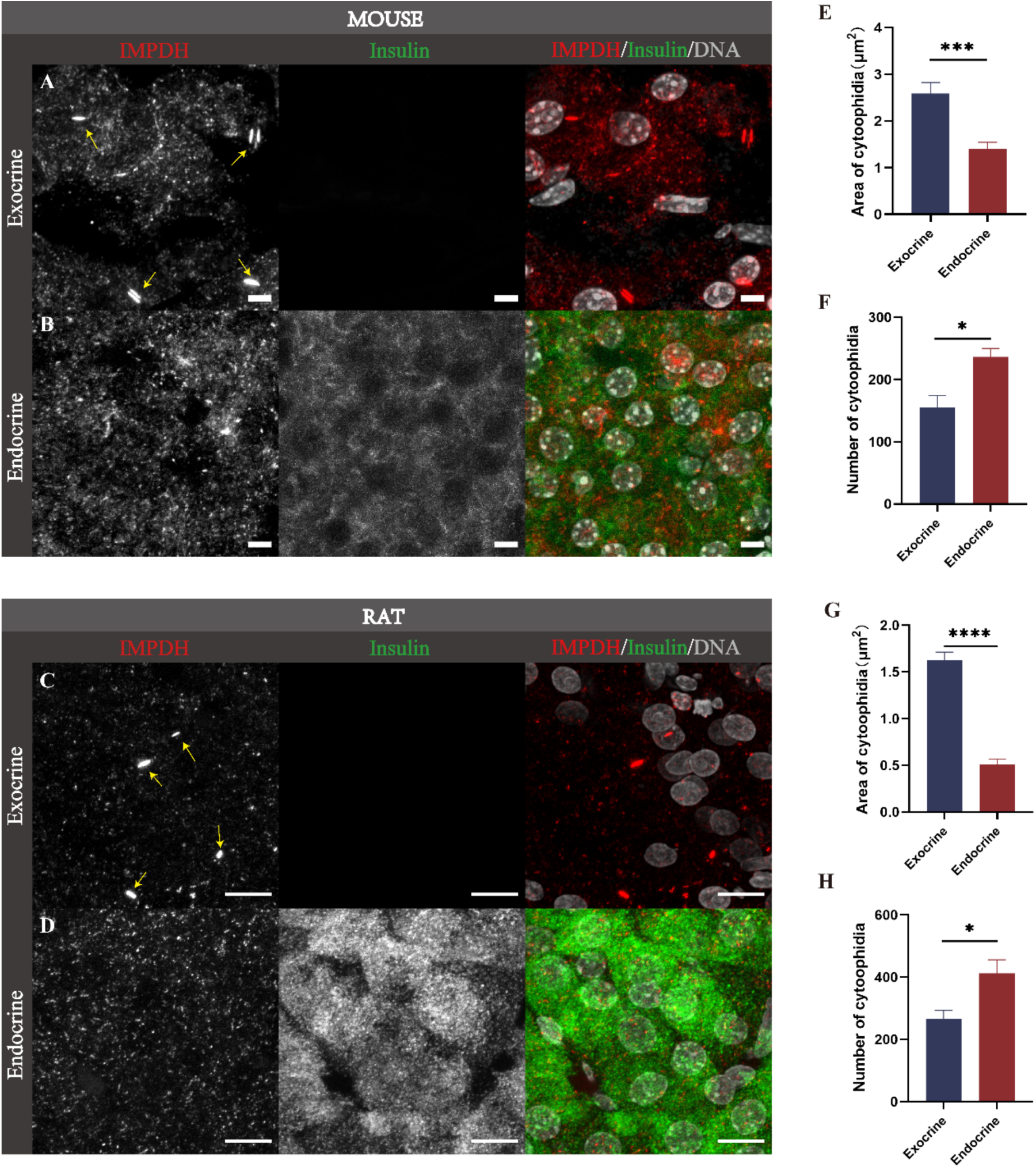
Differential distribution of IMPDH cytoophidia in islet and non- islet regions of the normal pancreas. **(A-B)** Immunofluorescence images of IMPDH cytoophidia in the pancreas of normal C57BL/6J mice (n=5). Scale bar: 10 μm. **(C-D)** Immunofluorescence images of IMPDH cytoophidia in the pancreas of normal SD rats (n=5). Scale bar: 10 μm. Panels (A-D) show single-channel grayscale images of IMPDH cytoophidia and merged images with IMPDH (red), insulin (green), and Hoechst 33342 (light gray). **(E-F)** Quantitative analysis of IMPDH cytoophidia area in normal mice. **(G-H)** Quantitative analysis of IMPDH cytoophidia area in normal rats. In (E-H), bar charts show the area of IMPDH cytoophidia in islet and non-islet regions. Statistical significance: *P<0.05, ***P<0.001, ****P<0.0001 (Mean ± SEM, T-test). Over 100 cytoophidia were analyzed per group.

### IMPDH Cytoophidia Elongate in Islets of Diabetic Models

To explore the relationship between IMPDH cytoophidia and islet function, we utilized diabetic models. Diabetes was induced in SD rats using streptozotocin (STZ), while BKS-db mice, which lack functional leptin receptors and spontaneously develop diabetes, served as an additional model. Immunofluorescence staining was performed to assess changes in IMPDH cytoophidia morphology under diabetic conditions. We labeled IMPDH cytoophidia with anti-IMPDH antibodies, islets with anti-insulin antibodies, and nuclei with Hoechst 33342. Hematoxylin and eosin (H&E) staining confirmed morphological alterations in diabetic islets, including irregular shapes and disrupted cell arrangements, validating the successful establishment of the diabetic model (Fig. 3A-D). Immunofluorescence analysis revealed a significant elongation of IMPDH cytoophidia within the islets of diabetic animals compared to controls (Fig. 3E-H). Quantitative measurements further supported this observation, demonstrating a marked increase in cytoophidia length in diabetic islets (Fig. 3I-J).

**Figure 3.**
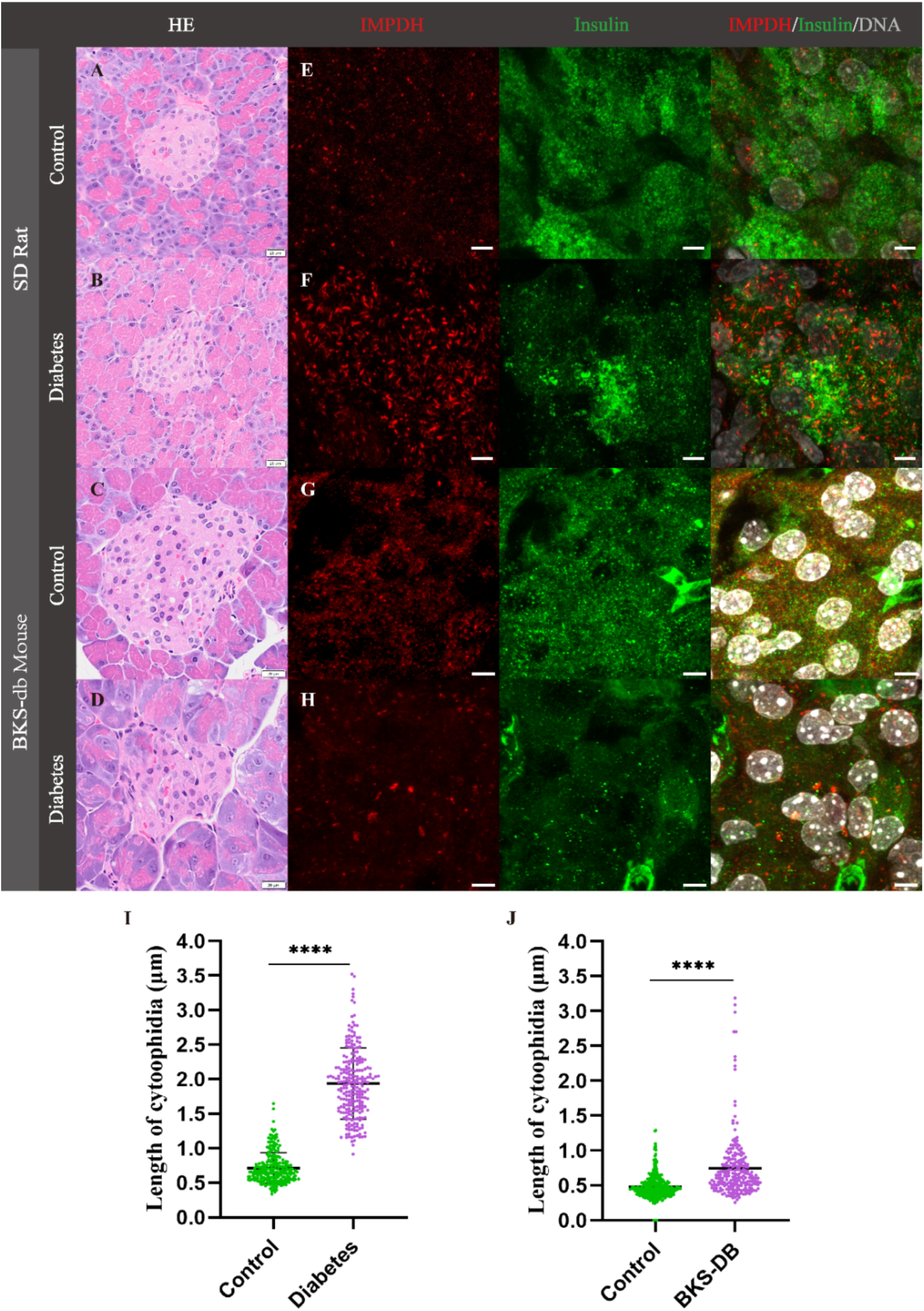
IMPDH cytoophidia elongate in islets of diabetic models. **(A-D)** H&E staining of pancreatic tissue. Scale bar: 20 μm. **(E)** Immunofluorescence image of IMPDH cytoophidia in islets of control SD rats (n=5). **(F)** Immunofluorescence image of IMPDH cytoophidia in islets of diabetic SD rats (n=5). **(G)** Immunofluorescence image of IMPDH cytoophidia in islets of control BKS- db mice (n=5). **(H)** Immunofluorescence image of IMPDH cytoophidia in islets of diabetic BKS- db mice (n=5). In (E-H), IMPDH is shown in red, insulin in green, and Hoechst 33342 (DNA, nuclei) in light gray. Scale bar: 5 μm. **(I)** Quantitative analysis of IMPDH cytoophidia length in islets of diabetic SD rats versus controls. **(J)** Quantitative analysis of IMPDH cytoophidia length in islets of diabetic BKS- db mice versus controls. Statistical significance: ****P<0.0001 (Mean ± SEM, T-test).

### Oleate Promotes the Assembly of Long IMPDH Cytoophidia

To investigate the potential mechanisms underlying cytoophidia elongation in diabetic conditions, we isolated primary islets from healthy mice for in vitro experiments. Given that circulating free fatty acids (FFAs), particularly oleic acid and palmitic acid, are elevated in diabetes, we focused on oleic acid due to its known role in promoting IMPDH translocation into lipid bodies. Primary islets were isolated and confirmed to retain intact morphology and insulin secretion capability via immunofluorescence staining (Fig. 4A) (32). When cultured with low concentrations of oleate (0.2 mM), no IMPDH cytoophidia were observed (Fig. 4B-C). However, at higher concentrations (0.4 mM and 1.6 mM), prominent elongated IMPDH cytoophidia formed within the islets (Fig. 4E-F), suggesting that elevated oleate levels may drive cytoophidia elongation in diabetic conditions.

**Figure 4.**
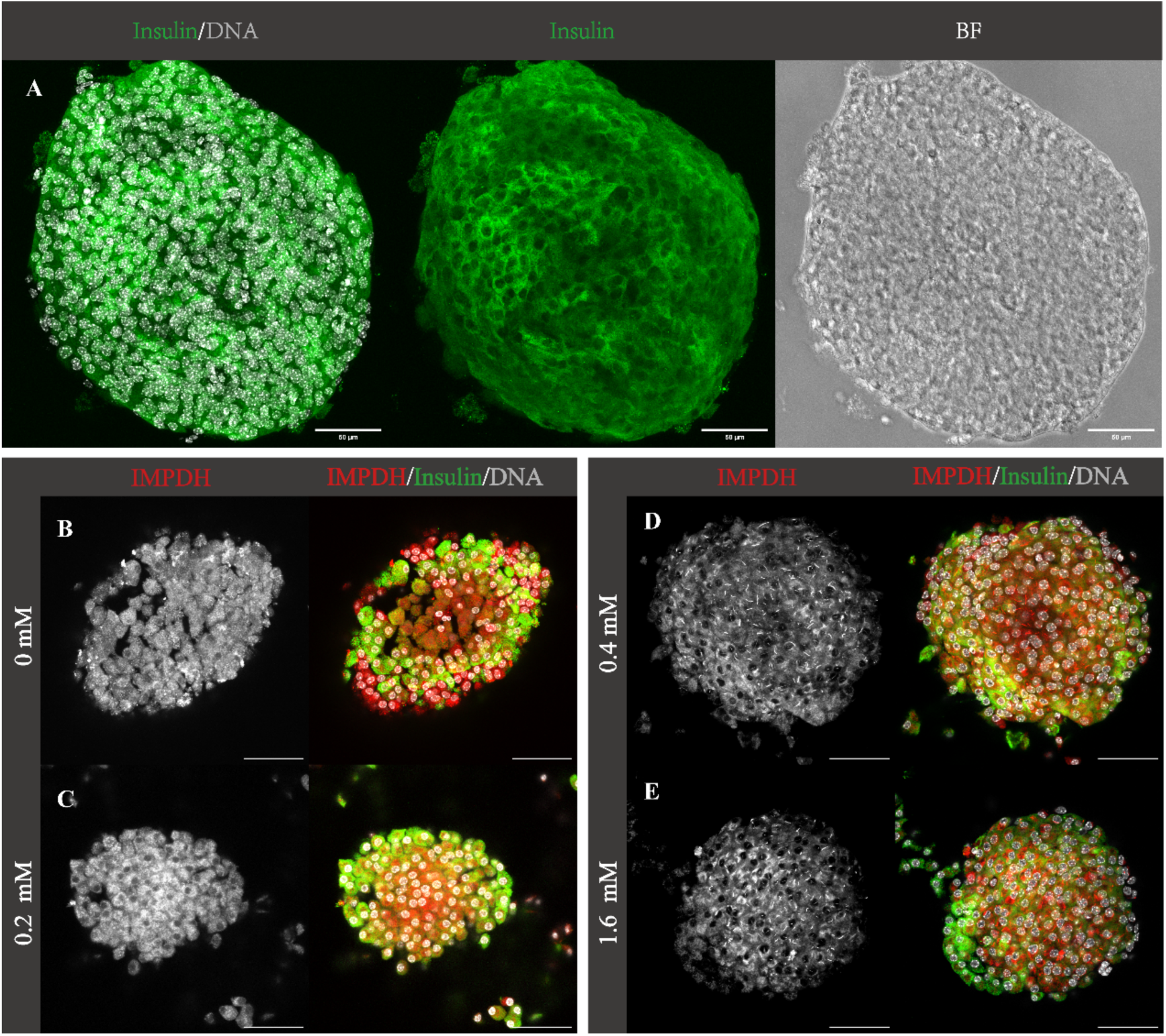
Oleate promotes the assembly of long IMPDH cytoophidia. **(A)** Immunofluorescence staining of isolated islets. Scale bar: 50 μm. **(B-E)** Immunofluorescence staining of IMPDH cytoophidia in isolated islets treated with oleate: **(B)** 0 mM oleate (control). **(C)** 0.2 mM oleate. **(D)** 0.4 mM oleate. **(E)** 1.6 mM oleate. Scale bar: 50 μm. Each group includes at least 50 primary islets. In (A-E), IMPDH is shown in red, insulin in green, and Hoechst 33342 (DNA, nuclei) in light gray.

### Elongation of IMPDH Cytoophidia Impairs Glucose Tolerance

To assess the functional consequences of cytoophidia elongation, we administered mycophenolic acid (MPA), a known inducer of cytoophidia assembly, to normal C57 mice. Pancreatic tissues were collected 10 hours post-injection, and immunofluorescence staining revealed significant elongation of IMPDH cytoophidia within the islets (Fig. 5A-C). Glucose tolerance tests (GTT) demonstrated that MPA-treated mice exhibited consistently higher blood glucose levels compared to controls from 10 to 45 minutes post-glucose administration, indicating impaired glucose metabolism (Fig. 5D). Analysis of the area under the curve (AUC) for GTT further confirmed reduced glucose tolerance in MPA-treated mice (Fig. 5E), supporting the hypothesis that cytoophidia elongation is associated with disrupted glucose homeostasis.

**Figure 5.**
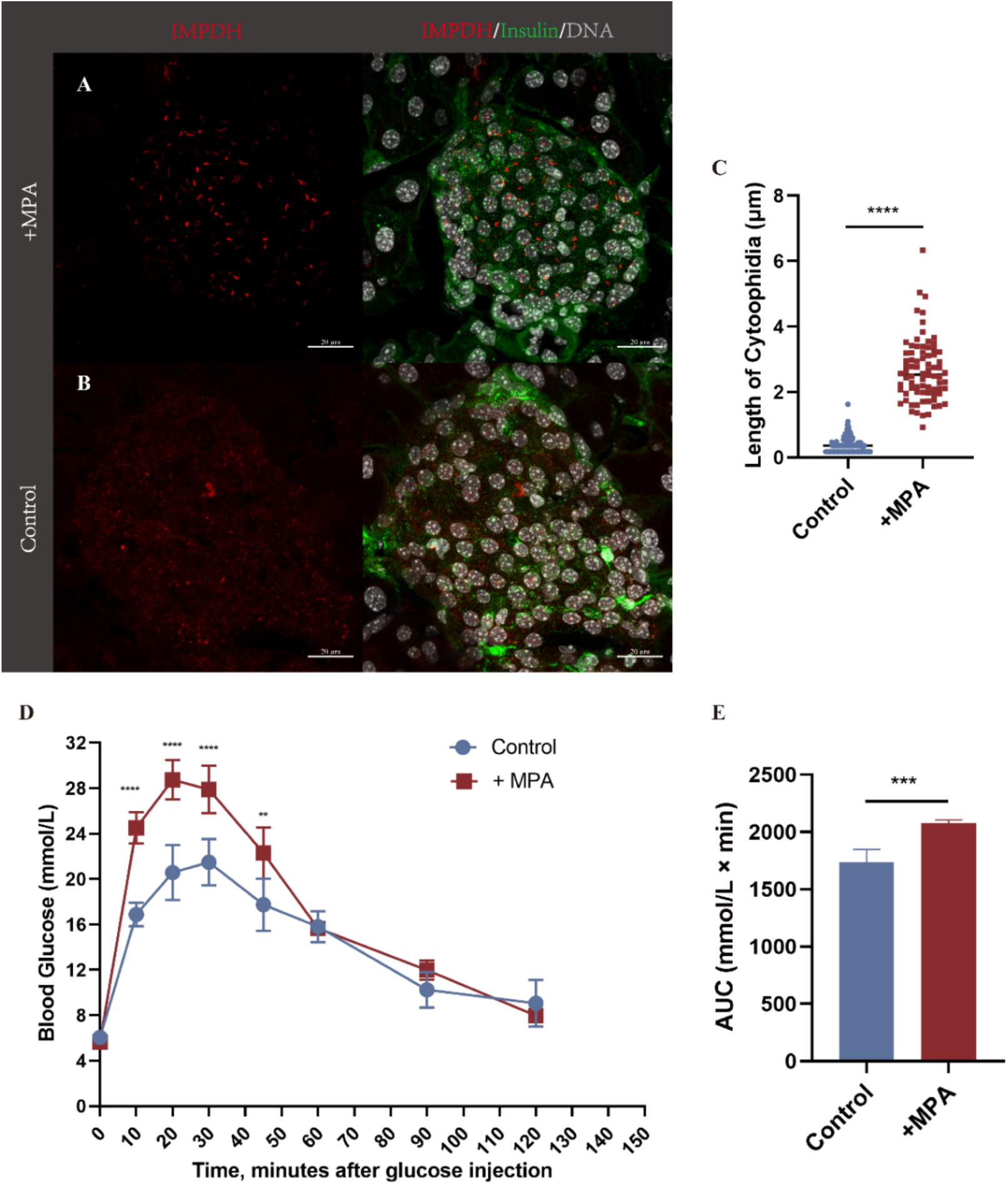
Elongation of IMPDH cytoophidia impairs glucose tolerance in mice. **(A-B)** Immunofluorescence images showing elongated IMPDH cytoophidia in islets of normal mice (n=5) 10 hours after MPA injection (500 mg/kg). Scale bar: 20 μm. IMPDH is shown in red, insulin in green, and Hoechst 33342 (DNA, nuclei) in light gray. **(C)** Quantitative analysis of IMPDH cytoophidia length in MPA-treated versus control mice. Over 100 cytoophidia were analyzed per group. **(D)** Glucose tolerance test (GTT) results showing blood glucose levels over time after glucose injection. **(E)** Area under the curve (AUC) analysis for blood glucose levels. In (C-E), control group (n=5) data are shown in blue, and MPA-treated group (n=5) data are shown in red. Statistical significance: ***P<0.001, ****P<0.0001 (Mean ± SEM, T-test).

## DISCUSSION

Our study provides novel insights into the role of IMPDH cytoophidia in the pancreas, revealing their distinct morphological distribution under both physiological and pathological conditions. We demonstrated that IMPDH cytoophidia are more abundant in islets compared to exocrine tissue and undergo significant elongation in diabetic models. Furthermore, we identified oleic acid and mycophenolic acid (MPA) as key factors promoting cytoophidia assembly. Notably, MPA-induced elongation of IMPDH cytoophidia was associated with impaired glucose tolerance, suggesting a potential link between cytoophidia dynamics and metabolic stress responses in islet cells. These findings highlight the importance of IMPDH cytoophidia in glucose metabolism and pancreatic function, offering new perspectives on their role in metabolic regulation.

### Expanding the Scope of IMPDH Cytoophidia Research

Previous studies on IMPDH cytoophidia have primarily focused on mice and human cells, with limited attention to other species. Our study is the first to report the presence of IMPDH cytoophidia in rat pancreatic tissues, thereby broadening the scope of research on these structures. Earlier work indicated that the number of IMPDH cytoophidia decreases in the pancreas of overnight-fasted mice, linking their assembly to nutritional status. In contrast, our findings reveal that cytoophidia elongation is associated with diabetic conditions, suggesting that their formation is also influenced by metabolic stress. This dual regulation by both nutritional and metabolic factors enriches our understanding of IMPDH cytoophidia dynamics and their potential role in cellular adaptation to stress.

### Mechanistic Insights and Future Directions

Our study utilized MPA, a well-known inducer of cytoophidia assembly, to investigate the functional consequences of cytoophidia elongation. While MPA treatment provided valuable insights, future studies could employ additional tools, such as the IMPDHY12A mutant, to further dissect the relationship between cytoophidia assembly and islet function. Moreover, our observation that oleic acid promotes cytoophidia elongation in vitro suggests a potential mechanism underlying cytoophidia dynamics in diabetic conditions. However, the downstream effects of oleic acid and other factors contributing to cytoophidia assembly in diabetes remain to be explored. Identifying these factors and their interplay with IMPDH cytoophidia will be critical for understanding their role in metabolic regulation.

### Limitations and Future Research

While our study provides significant advancements in the field, several limitations warrant consideration. First, our findings are primarily based on mouse and rat models, and future research should investigate the presence and behavior of IMPDH cytoophidia in other species, including clinical samples from diabetic patients. Second, although we identified oleic acid as a potential driver of cytoophidia elongation, the broader metabolic landscape of diabetes likely involves additional factors that remain to be elucidated. Third, while we observed a correlation between cytoophidia elongation and impaired glucose tolerance, the underlying mechanisms remain unclear. Future studies should aim to clarify how cytoophidia dynamics influence insulin secretion and pancreatic function at the molecular level.

## Conclusion

In summary, our study provides a comprehensive exploration of IMPDH cytoophidia morphology and distribution in the pancreas under different pathological conditions. By demonstrating the link between cytoophidia elongation and impaired glucose tolerance, we highlight their potential role in metabolic regulation and islet cell stress responses. These findings not only advance our understanding of IMPDH cytoophidia but also open new avenues for therapeutic strategies targeting metabolic-related diseases. Future research should focus on elucidating the molecular mechanisms underlying cytoophidia dynamics and their functional implications in metabolic health and disease.

## ACKNOWLEDGMENTS

We thank the Molecular Imaging Core Facility and Molecular and Cell Biology Core Facility at the School of Life Science and Technology, ShanghaiTech University for providing technical support. This work was supported by the grants from the Ministry of Science and Technology of China (No. 2021YFA0804700), Shanghai Frontiers Science Center for Biomacromolecules and Precision Medicine, Shanghai Ninth People’s Hospital, National Natural Science Foundation of China (Grant Nos. 32370744 and 32350710195), and UK Medical Research Council (Grant Nos. MC_UU_12021/3 and MC_U137788471).

## Notes

### Competing Interest Statement

The authors have declared no competing interest.

